# Who is who in necromass formation and stabilization in soil? Unraveling the role of fungi and bacteria as complementary players of biogeochemical functioning

**DOI:** 10.1101/2025.03.06.641408

**Authors:** Selina Lepori, Nadja Rohner, Xingqi Li, Xiaojuan Feng, Rota Wagai, Viviana Loaiza, David Sebag, Eric Verrechia, Daniel B. Nelson, Ansgar Kahmen, Claire Chenu, Pascal A. Niklaus, Anna-Liisa Laine, Luiz A. Domeignoz-Horta

**Affiliations:** Graduate Program in Quantitative Environmental Sciences, University of Zurich, Zurich, 8057, Switzerland; Department of Evolutionary Biology and Environmental Studies, University of Zurich, Zurich, 8057, Switzerland; State Key Laboratory of Vegetation and Environmental Change, Institute of Botany, Chinese Academy of Sciences, Beijing 100093, China; China National Botanical Garden, Beijing 100093, China; College of Resources and Environment, University of Chinese Academy of Sciences, Beijing 100049, China; Institute for Agro-Environmental Science, National Agriculture and Food Research Organization, Tsukuba, 305-0856, Japan; IFP Energies Nouvelles, Rueil-Malmaison, 92852, France; Faculty of Geosciences and the Environment, Institute of Earth Surface Dynamics, University of Lausanne, Lausanne, 1015, Switzerland; Department of Environmental Sciences – Botany, University of Basel, Basel, 4056, Switzerland; Université Paris-Saclay, INRAE, AgroParisTech, UMR EcoSys, 91120 Palaiseau, France; Research Centre for Ecological Change, Organismal and Evolutionary Biology Research Programme, Faculty of Biological and Environmental Sciences, University of Helsinki, Finland

**Keywords:** global change, microbial carbon use efficiency, necromass, carbon cycling, climate change, ecosystem resilience, soil organic matter

## Abstract

Multiple global change drivers have caused a large carbon (C) debt in our soils. To remedy this debt, understanding the role of microorganisms in soil C cycling is crucial to tackle the C soil loss. Microbial carbon use efficiency (CUE) is a parameter that captures the formation of microbially-derived soil organic matter (SOM). While it is known that biotic and abiotic drivers influence CUE, it remains unclear how distinct microbial communities and abiotic conditions influence the formation of microbially-derived SOC and its persistence in soils. Here, we combined the inoculation of distinct communities (a biotic factor) grown at different moisture levels (an abiotic factor) to manipulate the formation of microbial necromass in model soil. In a follow-up experiment, we then evaluated the persistence of this previously formed microbially-derived C to decomposition. While we show that necromass formation reflects the microbial community composition, the SOC formed within the most complex community of bacteria and fungi seems to be more resistant to decomposition compared to the SOC formed within the simpler communities (bacteria and fungi simple community, bacteria only and fungi only communities). Moreover, fungal necromass proved to be more thermally-stable than bacterial necromass, if this necromass is formed under the presence of bacteria. Our findings reveal that although abiotic factors can influence microbial physiology, the biological origin of microbially-derived C and the co-occurrence of fungal and bacterial growth were the stronger drivers explaining SOM persistence in these soils, suggesting the importance of microbial succession in SOC stabilization.

## Introduction

Soil organic matter (SOM) forms a significant portion of the C stored in terrestrial ecosystems^1^. In fact, soils serve as the largest carbon reservoir on land, housing 2000 billion tons of organic C^2^. This surpasses the carbon content found in vegetation and the atmosphere^3^, and it shows a considerably slow turnover rate^4^. However, SOM levels decline has led to an increase in CO_2_ emissions to the atmosphere. This decline is largely attributed to the conversion of natural land into agricultural soils^5^, intensification of agricultural practices^6^ and global warming^7^. Thus, it is crucial to better understand the drivers of SOM formation and persistence in soils.

Growing evidence suggests that SOM formation and stabilization occur through microbial processing^8^, making microbial biomass formation a key driver of SOM dynamics^9,10^. For example, microbially-derived necromass, which are the microbial products and residues that form a fraction of soil organic matter, were shown to be relevant to understand changes in SOM pools^9–12^. It has been estimated that up to 50% of the more stable SOM consists of microbial necromass^10,12^. Thus, SOM pools are regulated in part by the rate and efficiency with which soil microorganisms incorporate plant inputs into their biomass and thereby form more stable components of SOM^20^. A microbial parameter that allows studying microbial physiology capturing the fraction of carbon taken up by microbial cells and retained in biomass compared to the fraction being respired is carbon use efficiency (CUE)^15^. A previous study showed that microbial community structure influence community CUE^16^, suggesting that distinct communities can play a role in SOM formation rates. This same study showed that more diverse microbial communities allocate more C to growth in relation to respiration than species-poor communities^16^, which could indicate higher turnover rates of SOM under more complex communities. In a follow-up study it was observed that microbial activity and community composition drives the chemical fingerprint of newly formed SOM and its thermal-signature^17^. Previous studies have also showed that fungal and bacterial communities inhabiting soils exhibit distinct nutrient demands and differently influence the community CUE^18,19^ and these groups are considered to play different roles in soil C cycling^20^. In bacteria-only communities, the activity of extracellular enzymes is up to 75% lower than in communities containing fungi^17^. These results would suggest an overall smaller C uptake and reduced turnover of SOM in “bacteria-only” communities. Nevertheless, this study also showed that the SOM chemical signature was explained by the bacterial community composition, highlighting the need for further research on fungal and bacterial interactions and their contributions to the process of SOM stabilization. An underlying mechanism of fungal and bacterial interactions is cross-feeding, by which by-products of fungal metabolism can benefit bacterial communities leading to more efficient growth^21^. Cross-feeding occurs when the metabolic by-products excreted by one organism or population are used by others and are considered a key mediator of positive interactions within microbial communities. If microorganisms mobilize biomass building blocks from microbial necromass within the SOM^22^, cross-feeding might be also a mechanism enhancing SOM turnover. While previous results suggest that decomposition of fungal residues is a regulator for soil functioning and C accumulation in soils^23^, it becomes crucial to evaluate the role of fungal and bacterial interactions in this process.

Abiotic factors are known to directly influence microbial metabolism and by modifying the environment, they can also influence the biotic interactions^24^. For example, a study showed that drought can modulate the relationship between bacterial diversity and CUE, with CUE being positively correlated with bacterial diversity under high moisture but not under drought^16^. Drought is also known to have different effects on soil fungal and bacterial communities^25^ and their growth efficiency^26,27^. Soil bacterial communities are expected to be more affected by drought as they are more susceptible to substrate limitation^28^, desiccation and/or water deficit compared to fungal growth which is less dependent on the water film due to hyphal growth^29^. Abiotic factors such as wet/dry cycles will also impact soil aggregate structure and formation^30^. For example, fungal hyphae was showed to be able to enmesh soil particles even under drier conditions^31^. Soil aggregation is an important parameter contributing to the persistence of SOM in soils^32^.

The study of the formation of more stable components of SOC and their turnover, despite challenging^33^, is a growing research area^34^. More recently, SOC ramped thermal analysis have been suggested as a way to capture carbon pools differing in their residence-time and ”biologically-reactivity”^35^, as it provides a metric of SOM intrinsic thermal stability^36,37^. To gain further insights into the roles of fungal and bacterial communities for C cycling through soils we designed an experiment where we manipulated microbial communities and assessed its impact on CUE, necromass and aggregate formation as well as the thermal stability of the formed SOM. The experiment consisted of two phases: in phase I, we inoculated a sterile C-free soil with four distinct communities (complex community of fungi and bacteria, simple community of fungal and bacterial strains, bacteria only and fungi only communities). For phase II, we then X-ray sterilized the soils, re-inoculated them, and evaluated how the new microbial community degraded the microbially-derived SOC formed during the phase I (Figure 1). We hypothesized that: 1) Fungi-dominated community produce a higher content of necromass compared to bacteria-dominated community; 2) more complex microbial communities result in a microbial-derived SOC that is more persistent in the soil due to enhanced SOC turnover; and 3) Co-occurrence of fungal and bacterial growth enhances SOM thermal-stability promoting potential SOC stabilization.

**Figure 1.**
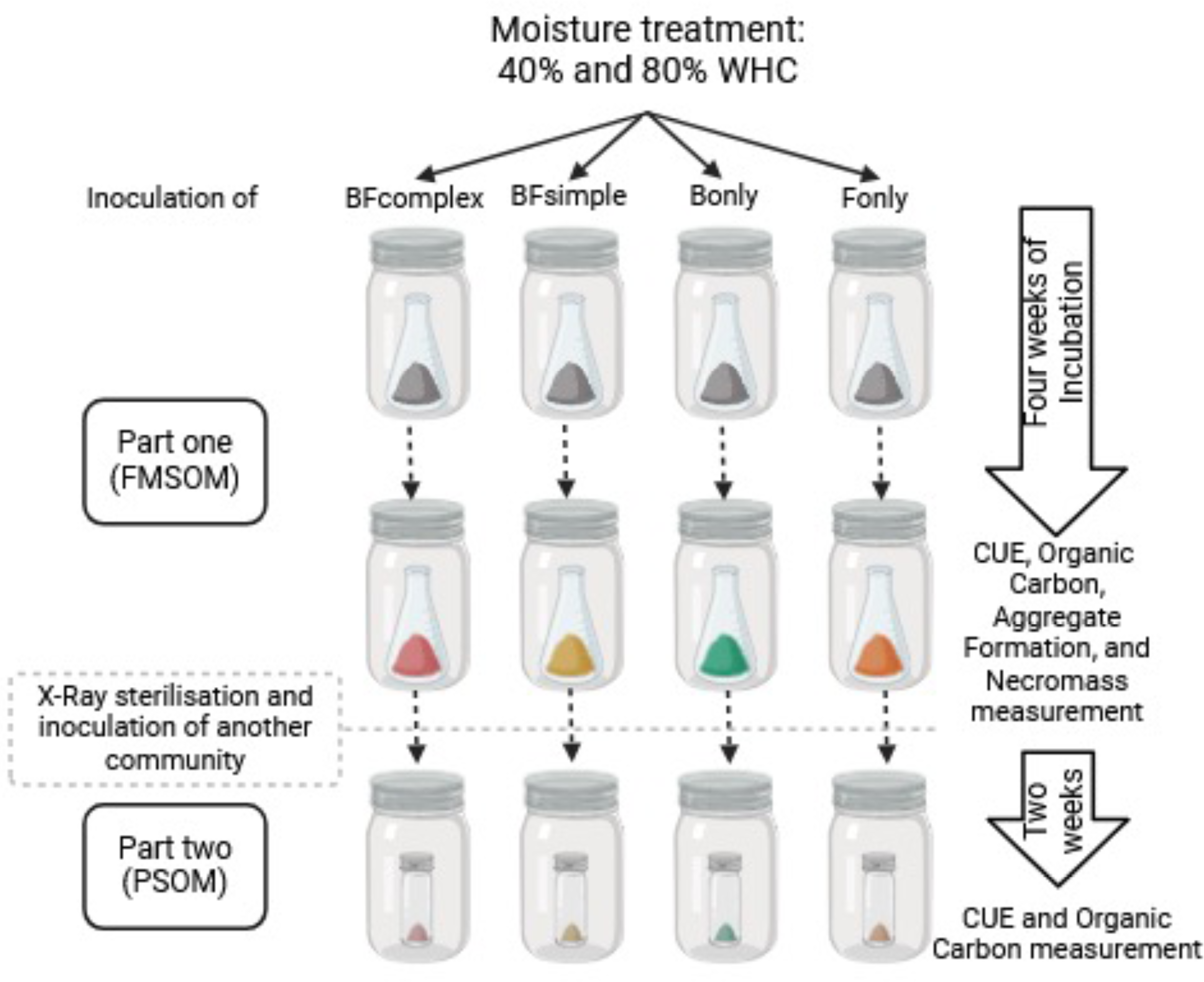
Experimental design. Distinct communities with equivalent densities were inoculated in an initially microbe-free and C-free soil. The inoculum was manipulated by extracting a complex bacteria and fungal community from a natural soil (“BF_complex_”); inoculating two bacterial strains (“B_only_”) (representing an Actinobacteria and Betaproteobacteria: *Streptomyces sp* and *Microvirgula aerodenitrificans, respectively*); inoculating one fungal strain (“F_only_”) (representing Ascomycota: Trichoderma koningii); combining the B_only_ and F_only_ inoculums into a “BF_simple_” inocula. These communities were incubated during four weeks in a phase referred as Formation of microbially-derived soil organic matter (“FMSOM”). We measured CUE, quantified the C remaining and its thermal signature as well as the necromass formed during the incubation and the soil aggregate formation (via physical fractionation). At the end of FMSOM all communities were X-ray sterilized. A new community derived from a natural soil was inoculated to estimate the persistence of the microbially-derived carbon formed during the first weeks of incubation (Persistence of Soil Organic Matter; PSOM) and we performed this by measuring the respiration, growth and CUE at this second incubation phase.

## Materials and Methods

### Model soil, inoculation and incubation conditions

We created a microbe and C-free soil to evaluate how distinct microbial communities generate distinct microbial-derived-SOC. 90% acid-washed sand (0.1 - 0.8 mm) was mixed with 10% bentonite clay (montmorillonite) treated with calcium chloride and autoclaved three times with a minimum interval of 48hrs between cycles.

We created four different microbial community treatments. First, a simple community of bacteria (“B_simple_”) was created consisting of the mixture of two bacterial strains (*Streptomyces sp.* (gram-positive Actinomycetota DSM 687) and *Microvirgula aerodenitrificans* (gram-negative Pseudomonadota DSM 736). Then, a simple fungal (“F_simple_”) inocula treatment was generated using the fungal strain *Trichoderma koningii* (DSM 63059). To create a simple bacterial and fungal communities (“BF_simple_”) we used half of the inocula used to prepare the communities mentioned previously. Finally, we also created a complex community of fungi and bacteria (“BF_complex_”) by extracting an inocula from an agricultural soil from Finland originated from the TwinWin experiment^38^ and by shaking 2 g soil in 50 ml of sterile deionized water (Supplementary table 1). We aimed to inoculate the model soil with the distinct communities at the same abundance per g soil, and for this we determined the corresponding colony forming units (CFU) at different optical densities during the growth of the strains to determine the volume of inocula needed to reach the same abundance of about 1.0 x 10^7^ CFU g^-1^ soil. For the BF_complex_ community we performed serial dilutions and CFU countings to also determine the ratio of inocula:soil needed to reach about 1.0 x 10^7^ CFU g^-1^ soil. We also had non-inoculated microcosms which did not receive an inoculum (uninoculated controls; 8 samples) which we used to verify that these soils were sterile in the beginning of the experiment by evaluating CO_2_ production in these microcosms. Microcosms were incubated at two different water content treatments (40 and 80% WHC) in a full factorial design with four replicates per treatment (4 community treatments x 2 moisture levels x 4 replicates = 32 biological samples) (Figure 1). To allow for maximum utilization of nutrients, only one substrate addition was performed at the moment of inoculation. Each 10 g model soil microcosm was amended with Tryptic soy broth medium and glucose to result in 4.8 mg C g^-1^ soil.

### Formation of microbially-derived SOM (FMSOM)

We used an array of complementary methods to characterize the physiology of microbial communities across the distinct treatments, and the quantity and quality of microbially-derived C at the end of the FMSOM. Below, we describe in more detail the distinct methods. Microcosms were incubated for four weeks and microbial activity monitored by CO_2_ flux measurements every third day. This first phase of the experiment is denominated “Formation of Microbially-derived SOM” (FMSOM). At the end of the FMSOM incubation phase we determined the CUE of the communities with the ^18^O-H_2_O method^39,40^. We evaluated the quantity and quality of SOC with the ramped thermal rock-eval pyrolysis method^36^, measured the amino sugars produced by the distinct microbial communities during the incubation following established methods^41^, and estimated the aggregate formation by density fractionation in the different soils^42^.

### Cumulative respiration

The microcosms were placed inside a jar (0.75 l) containing a septum port that allowed sampling of the headspace and monitoring CO_2_ produced during microbial respiration. The CO_2_ respiration from these jars was measured by sampling the headspace 24 and 48 hr after closing the jars. After the second sampling, the jars were opened for a few minutes to reduce CO_2_ levels and to prevent microcosms from becoming anaerobic. Jars without any microcosms were also closed in order to measure the starting CO_2_ levels. The CO_2_ measurements were performed consecutively during the 4 weeks of incubation to allow us to calculate the total cumulative respiration from each microcosm and treatment. The CO_2_ concentration inside the jars was measured by gas chromatography – flame ionization detector (Agilent Technologies 7890A, CO_2_ was first reduced with H_2_ using a Nickel catalyst and then detected using a flame-ionisation detector).

### Growth, respiration, and CUE measurements

We used the substrate-independent method H_2_^18^O-CUE to evaluate the community CUE^39^. Weighing for the respiration and growth measurements started the day following the soil harvest after 4 weeks of incubation. Two 0.3 g soil aliquots were weighed into 15 ml falcon tubes and placed in a large tube. We controlled for soil moisture loss by regularly weighing the tubes. CUE was measured at the respective WHC treatments (i.e. at 40% and 80% WHC) and soil samples received 20% of the total water present as H_2_^18^O. Control soil samples received water with natural ^18^O abundance to subtract the natural abundance of ^18^O when calculating the ^18^O enrichment of the DNA. After adding water, the tubes were sealed and incubated at room temperature for 48 hours. Following this incubation period, 25 ml of the headspace was sampled for CO2 measurements. The soil samples were then frozen at -80^0^C until DNA extraction. CUE measurements using the H_2_^18^O-CUE method estimate the new microbial biomass produced during the incubation period based on ^18^O-DNA at the end of the incubation period. DNA was extracted from all soils incubated with ^18^O-water and a subset of soils incubated with ^16^O-water using the Qiagen Power Soil kit. Technical duplicates were pooled before quantification using Qubit. DNA δ^18^O values were measured using a Flash IRMS elemental analyzer operated in pyrolysis mode coupled to a Delta V Plus isotope ratio mass spectrometer (Thermo Fisher, Waltham, MA, 410 USA) at the Stable Isotope Ecology Laboratory, University of Basel, Switzerland, and results are reported relative to Vienna Standard Mean Ocean Water (VSMOW) in ‰. Long term instrumental precision for the laboratory for non-^18^O-enriched analyses is 0.2 ‰. Analytical precision for ^18^O-enriched samples measured as a part of this study was 1.57 ‰ (n = 6). CUE was calculated using a conversion factor Microbial Biomass Carbon:DNA of 10.9 according to Spohn^39^.

### SOM quantity and quality

We used ramped thermal rock-eval pyrolysis to evaluate SOM quality and quantity^37^. During rock-eval, carbon dioxide is quantified as it comes off a soil sample subject to increasing temperatures, thereby providing a metric of SOC intrinsic thermal stability. Compounds with high thermal stability include aromatic and phenolic non-lignin compounds, while lipids and polysaccharides tend to have lower thermal stability^35^. Soils were dried at 65°C and crushed to a fine powder in a mortar and pestle. Between 50 and 70 mg soil was pyrolyzed over a temperature ramp from 200 to 650°C, followed by combustion to 850°C using a rock-eval 6 pyrolyzer (Vinci technologies) at the Institute of Earth Sciences of the University of Lausanne (Switzerland). Hydrocarbons released during the ramped pyrolysis process were measured by a flame ionization detector (FID). Rock-eval was performed in the soils at the end of the FMSOM phase and also at the end of the second incubation phase (further explained below). To evaluate if distinct amount of easily available dissolved hydrocarbons (i.e. amended glucose) remained in the different treatments at the end of FMSOM we used two approaches: 1) we estimated the remaining hydrocarbons considering the cumulative C-losses via respiration in addition to the necromass formed during the incubation and microbial biomass as previously; and 2) we measured ultraviolet absorbance at 254 nm as a proxy of remaining dissolved organic carbon. For this, 2 g of soil was mixed with 8 ml deionized water and vortexed for 10 seconds. After 30 minutes of settling, 4.5 ml of the supernatant was extracted with a 5 ml syringe and transferred to another falcon tube through a filter (PES, Merk Millipore Ltd., Ireland). Then, 1.5 ml were pipetted into a quartz cuvette and the optical density (OD) was measured with a SpectraMax M2 plate reader using the deionized water as blank.

### Microbial necromass (amino sugars) produced during incubation

Microbial necromass was extracted from freeze-dried soil (1 g aliquot) in duplicates^44^. Then, soils were hydrolyzed (6 M HCl, 105°C, 8 h). The precipitates were removed and hydrolysates adjusted to a pH of 6.6–6.8. Then, amino sugars were dissolved in methanol and separated from salts by centrifugation. After the addition of a quantitative standard (methyl-glucamine), amino sugars were transformed into aldononitrile derivatives^45^. The derivatives were further acetylated with acetic anhydride. The amino sugars were quantified on a Trace GC 1310 gas chromatograph coupled to an ISQ mass spectrometer (Thermo Fisher Scientific, USA) using a DB-5MS column (30 m × 0.25 mm i.d., film thickness, 0.25 μm) for separation. We used myo-inositol as a recovery standard. The recovery of amino sugars was in the range of 95%. We evaluated the production of glucosamine, mannosamine, galactosamine and muramic acid. Galactosamine and muramic acid are expected to occur predominantly in bacteria while glucosamine is predominantly produced by fungal cells. We calculated the amino-sugar-C equivalent expected to be of fungal or bacterial origin based on Joergensen^46^ considering that muramic acid (MurN) occurs only in the bacterial cell wall and glucosamine (GlcN) occurs mainly in fungal cell walls. We also calculated the amino sugar accumulation efficiency (AAE) using the following formula: AAE = (AminoSugar-C / (AminoSugar-C + cumulative(CO_2_-C)) * 100. AAE is reported in percent (%).

### Density fractionation

We used density fractionation approach to assess if microbial community activity promotes organo-mineral aggregation after the FMSOM incubation phase. To distinguish the organo-mineral aggregates from the initial mineral and organic particles, we isolated meso-density fraction (1.8 - 2.4 g cm^-3^) because the particle density of sand and bentonite clay was >2.4 g cm^-3^ (a pilot test showed 0.47% of the initial mineral mass had the particle density < 2.4 g/cc) and that of any organic compounds are <1.8 g cm^-3^ ^42^. Sodium polytungstate (SPT) was used to adjust solution density (SPT-0 grade, TC-Tungsten Compounds, Germany). Following a previous protocol^40^, freeze-dried samples (5 gram) was mechanical shaken for 30 min with SPT solution adjusted to 1.8 g cm^-3^ to disrupt the less-stable aggregates and isolated organic matter not associated with minerals as low-density fraction (<1.8 g cm^-3^) by recovering floated materials after centrifugation (2330 g). To obtain the meso-density fraction, the remaining liquid was replaced with 2.4 g cm^3^ SPT solution, shaken and centrifuged following by the recovery of the floatable as previously done. The remaining material was recovered as high-density fraction. The recovered low-density fraction was rinsed with deionized water using a vacuum filtration system with 0.45 μm membrane filter. After recovering meso- and high-density fractions, the material was transferred to 250 ml bottles and mixed with deionized water and centrifuged (17.000 g) to remove the salt until the salt concentration in the supernatant was below 50 uS l^-1^. The fully-rinsed fractions were freeze-dried for chemical analysis. For the three replicates from the four treatments (BF complex dry, wet, B only moist, F only moist) fractionated, the mass recovery was 94.1 % ± 1.0 % (mean ± SD).

### Persistence of microbially-derived SOM (PSOM)

After the first phase of the experiment (described above) in which we evaluated how distinct communities promoted the formation of microbial-derived soil organic matter (FMSOM), we sterilized the soil with X-ray radiation (40 kGy). We decided to sterilize with X-ray to avoid modification of recently formed SOM with more traditional methods such as autoclaving the soil. These microcosms were then inoculated with a complex inoculum similar to our BF_complex_ treatment obtained from the same agricultural experimental site (Fig. 1). The soils were incubated for 15 days to evaluate how the new inoculum would consume the recently formed microbial-derived SOC. To achieve this, we measured cumulative respiration during the incubation and at the end of this period we evaluated the microbial community CUE with the H_2_^18^O-CUE method and the quantity and quality of the SOC at the end of this incubation phase with the RE pyrolysis.

### Statistical analysis

All figures and statistical models were performed in R (Version 2023.09.0+463) using the packages *vegan*^39^, *agricolae*^48^, and *ggplot2*^49^. The response variables were log-transformed to fit model assumptions. Amino sugars data were normalized by dividing by the remaining total organic carbon in a sample and are expressed as amino-sugar-C per g SOC. The effects of inoculum and moisture treatments were tested by analysis of variance (ANOVA) and *post hoc* Tukey HSD tests. Linear models were used to assess the relationship between the amino-sugar-C produced during the first phase of the experiment and the microbial physiology measurements of the second part of the experiment. To evaluate how the distinct amino sugar residues in the soil relates to the SOC thermal signature measured during the pyrolysis phase of Rock-eval we performed spearman correlations between the distinct measured amino-sugars and the FID signal captured at each temperature by ramped Rock-eval analysis. Significant explanatory variables for respiration, growth and CUE were chosen by linear regression, model selection (backward) and by minimizing the Akaike Information Criterion (AIC). The statistical significance was assessed by 1000 permutations of the reduced model. A variance partitioning approach was then used to evaluate the relative contribution of those variables to explain the variation in respiration, growth and CUE measured in the PSOC phase using the function *varpart*^50^.

## Results

### Microbial activity and formation of microbial-derived residues during FMSOC

To assess the activity of the microbial communities we quantified cumulative respiration during the 30 days of incubation (Fig 2a-b). Cumulative respiration did not differ among communities except for the F_only_ treatment in which respiration was lower compared to BF_complex_ and BF_simple_ (F_3,27_ = 4.770, *p* = 0.007). Cumulative respiration was unaffected by the moisture treatments (F_1,27_ = 0.194, *p* = 0.663). The compilation of respiration measurements during the 30 days of incubation suggests no major difference in microbial activity across treatments except for the Fonly treatment.

**Figure 2.**
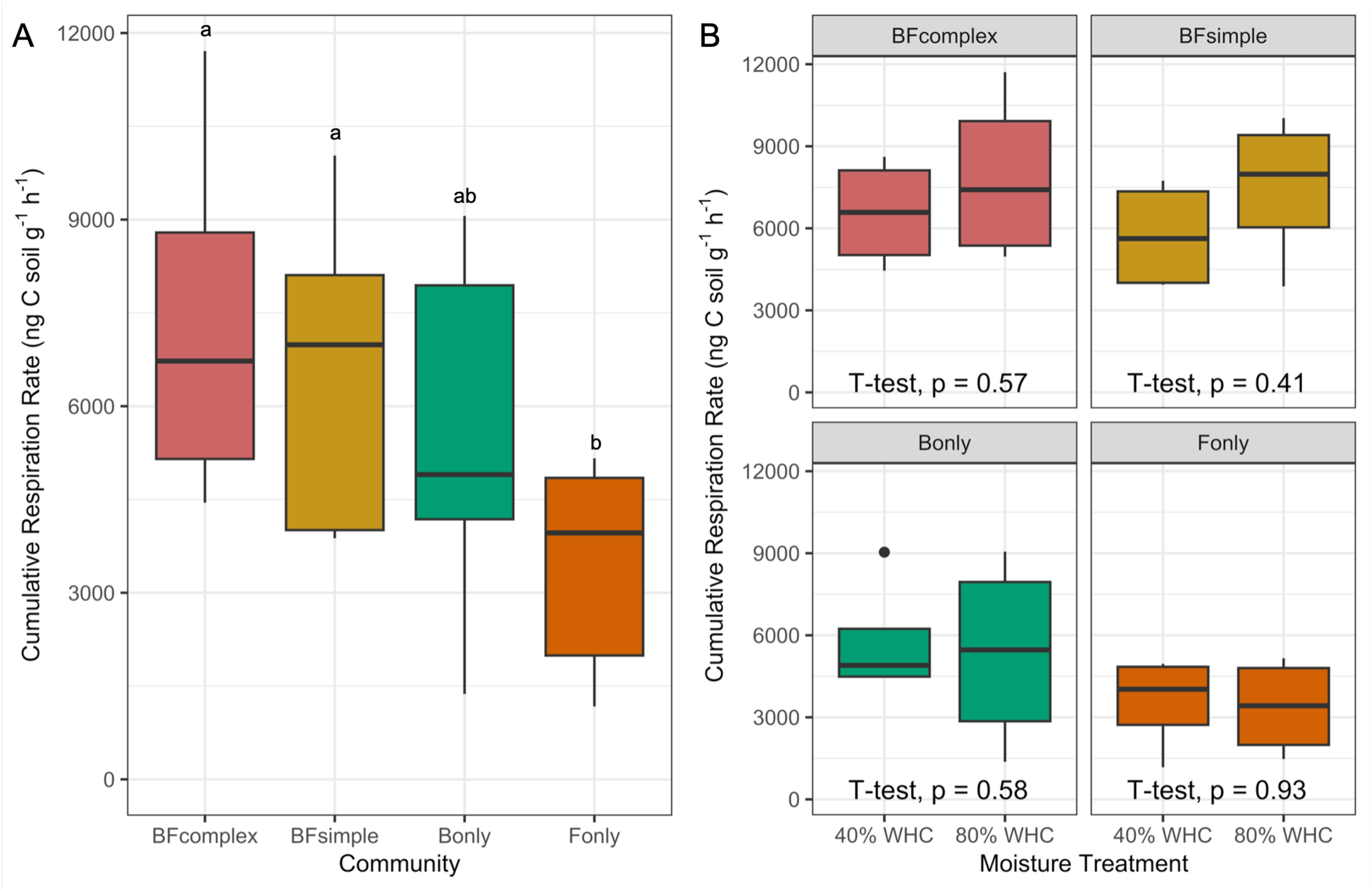
Respiration rates as a proxy for microbial activity. Boxplots showing the median, the interquartile range, the minimum and maximum values. Letters indicate significant differences between communities and moisture treatment. Cumulative respiration measured during first incubation phase (FMSOC) at the distinct communities (log transformed) (A). Cumulative respiration at each community type and moisture treatment (B). Significant differences between treatments are indicated with different letters (anova followed by Tukey HSD test, P < 0.05) or significant t-test results (P < 0.05).

Microbial necromass was measured to assess the influence of different microbial communities on SOC formation, using amino sugars formed during the 30-day incubation period as a proxy (Figure 3). The amino sugar signature reflected the microbial composition in the distinct soil samples (F_3,24_ = 3.778, *p* = 0.022). Moisture levels and the interaction of the community with moisture showed no significant effect (F_1,24_ = 1.728, *p* = 0.201, and F_3,24_ = 0.784, *p* = 0.514). Predominantly produced by fungi, glucosamine values were higher in the fungi-dominated soils (F_only_) than in the bacteria-dominated communities (B_only_) (Fig. 3A, 3F), while galactosamine and muramic acid, which are mainly produced by bacteria, were below detection limit in the fungi-dominated microcosms (Fig. 3C, 3D, 3E). We calculated the amino-sugars-C expected to be of fungal or bacterial origin based on Joergensen et al.^46^ (see methods). While the production of specific amino-sugars differs between communities (Fig. 3A-D), the total amino-sugar production was not different among communities or moisture treatments (Fig. 3F). The Amino-sugar Accumulation Efficiency (AAE), defined as the total amino-sugar formed in the soils relative to the total respiration, ranged from 0.60 % (CI_95%_= [0.44 - 0.75]) in the BF_complex_ up to 2.66% (CI_95%_= [0.88 - 4.44]) in the F_only_ treatment on average (F_3,28_ = 7.813, *p* < 0.05) (Figure 4).

**Figure 3.**
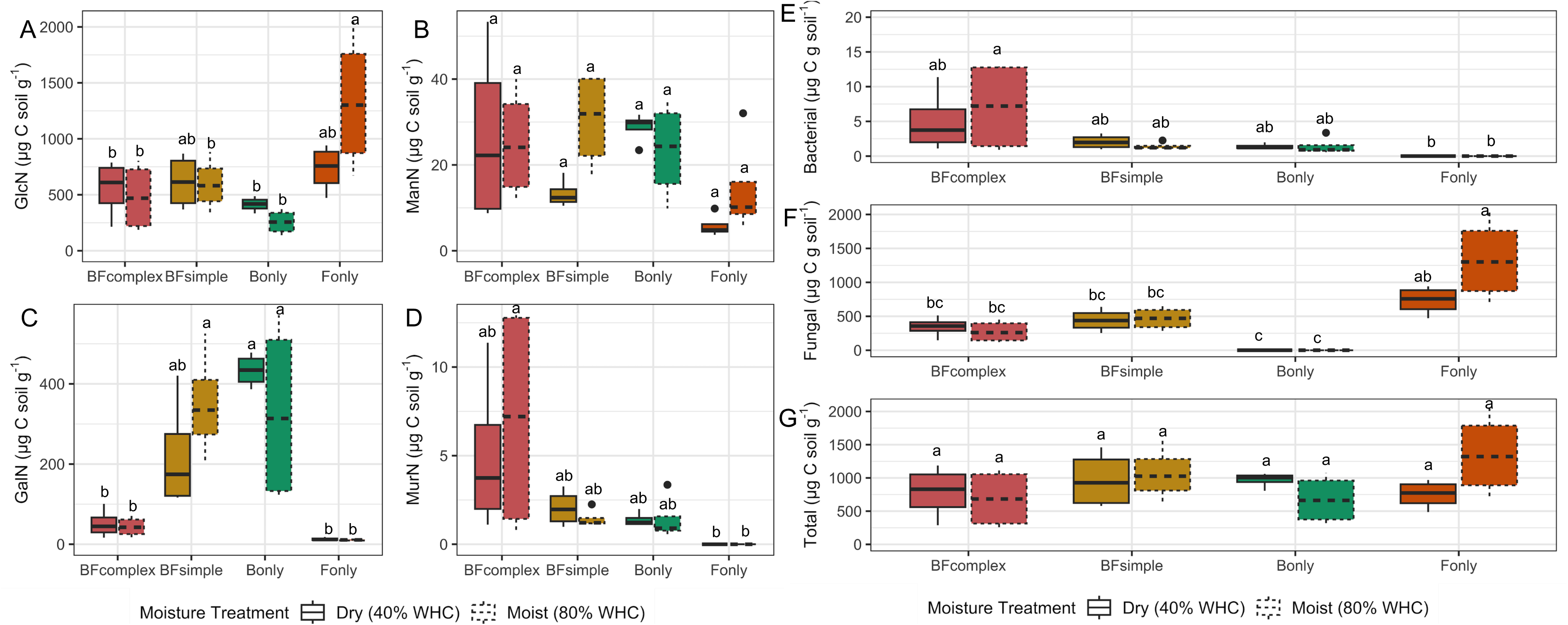
Amino sugar accumulated during FMSOM by each microbial community at two moisture treatments. Boxplots showing the median, the interquartile range, the minimum and maximum values. Letters indicate significant differences between communities and moisture treatment. Glucosamine (A), mannosamine (B), galactosamine (C) and muramic acid (D) produced during 30 days of incubation. Amino sugar of bacterial origin (E), fungal origin (F) and total amino sugar (G). Significant differences between treatments are indicated with different letters (anova followed by Tukey HSD test, P < 0.05).

**Figure 4.**
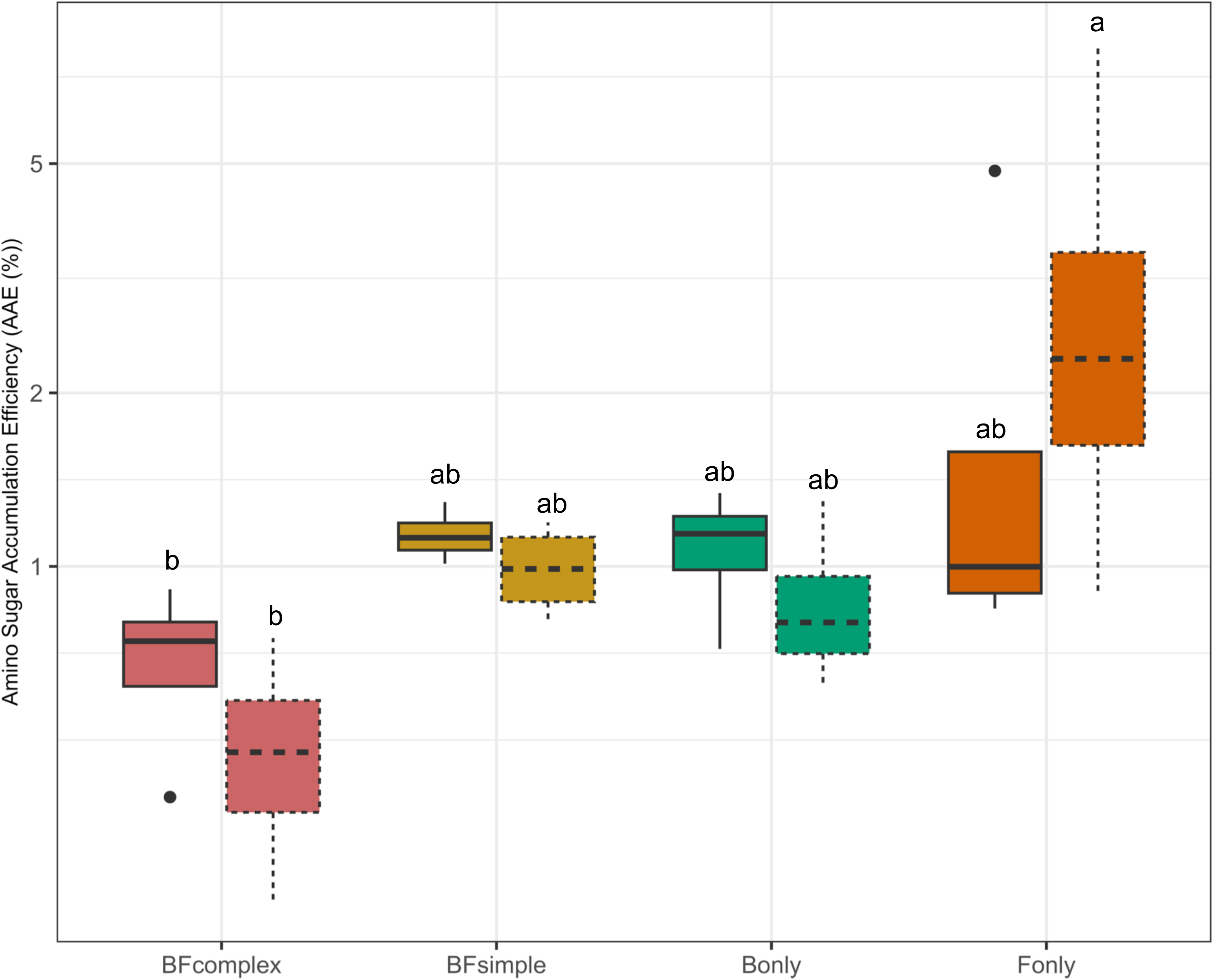
Amino Sugar Accumulation Efficiency (AAE; %). Boxplots showing the median, the interquartile range, the minimum and maximum values. Letters indicate significant differences between communities. Amino sugar-C accumulated at the end of the FMSOC after four weeks of incubation taking into account the cumulative respiration produced by each specific microcosm.

At the end of the FMSOM incubation phase we measured the growth with the H_2_^18^O-method and also respiration in a subsample of soil. While we observed some differences among communities for respiration and growth no significant difference was observed within a community for the distinct moisture levels (Fig. 5A, B). Respiration rates ranged from 795 ng CO_2_-C g^-1^ soil h^-1^ (CI_95%_= [180 - 1411]) for BF_complex_ to 8140 ng CO_2_-C g^-1^ soil h^-1^ for B_only_ (CI_95%_= [3296 - 12984]) while growth rates ranged from 17 ng C g^-1^ soil h^-1^ (CI_95%_= [11 - 22]) for BF_simple_ to 104 ng C g^-1^ soil h^-1^ (CI_95%_= [39 - 169]) for B_only_. Soil moisture did not affect respiration or growth significantly within a community. However, respiration and growth showed the opposite tendency. While respiration rates tended to be higher under the low moisture compared to high moisture treatment, growth was on average higher in moist soils compared to the dry soils (for growth: t = 2.80, df = 19.50, p-value = 0.01). The exception was the F_only_ treatment which showed the same pattern for both respiration and growth measurements. CUE is a compilation of growth and respiration measurements, as a consequence of these distinct responses we observed a significantly lower CUE in the dry BF_complex_ soils compared to the BF_complex_ wet soils (Fig. 5C). This same trend was observed for the CUE of BF_simple_ and B_only_ but the differences were not significant (Fig. 5C). F_only_ showed similar median values for CUE in both moisture treatments (average BF_simple_ = 2.0 %, average BF_complex_ = 12.8 %, CI_95% BFsimple_= [1.0 - 2.9], CI_95% BFcomplex_= [5.2 - 20.4]).

**Figure 5.**
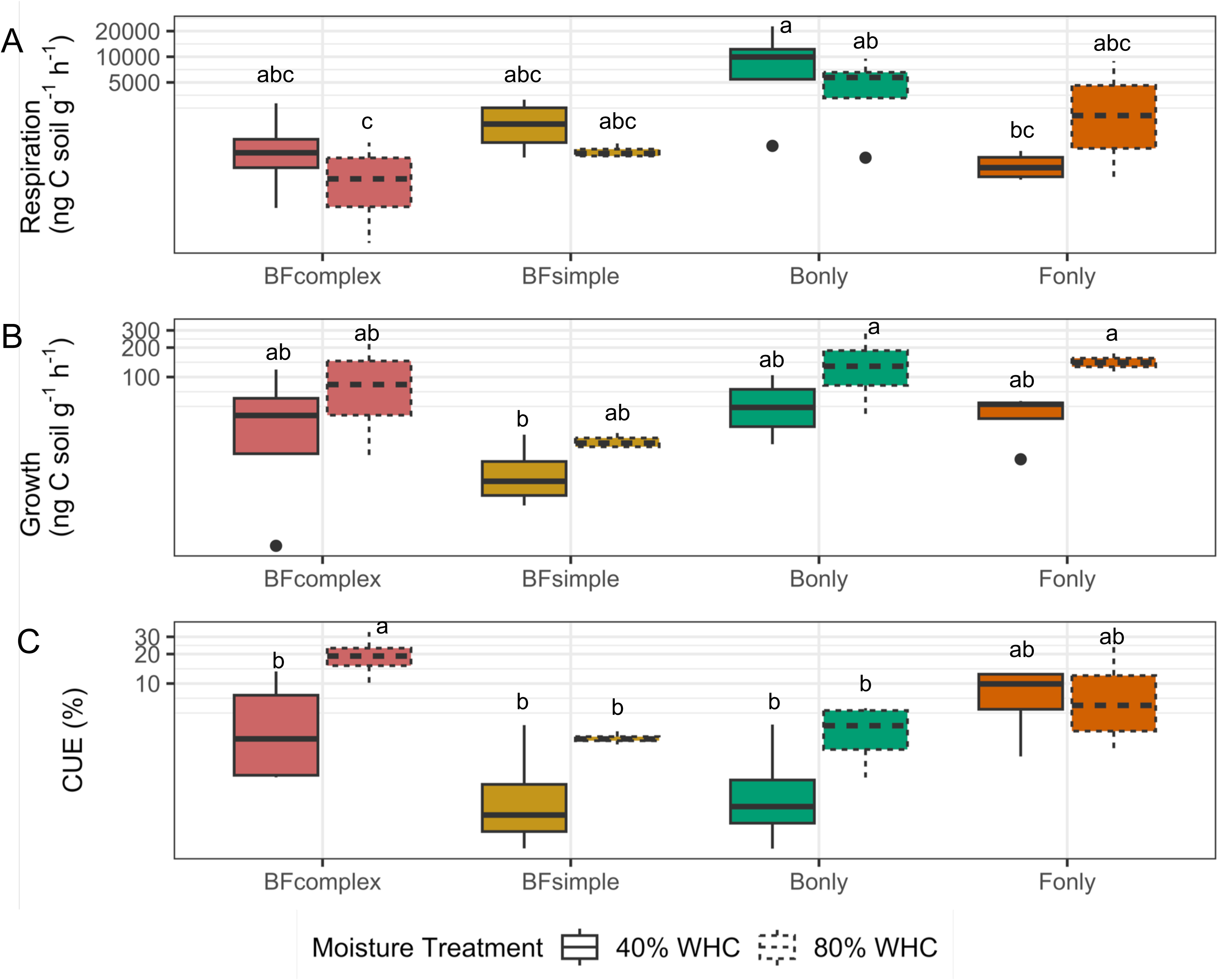
Carbon cycling processes measured at the end of the FMSOM incubation phase. Boxplots showing the median, the interquartile range, the minimum and maximum values. Letters indicate significant differences between communities and moisture treatment. Respiration (A), growth (B) and CUE (C) measured at the distinct communities under different moisture levels.

### Microorganisms’ influence on SOM formation and its thermal stability

The total organic carbon (TOC) did not differ among treatments at the end of the first incubation phase (Supplementary Figure 1a) suggesting that the microbial communities consumed the substrate added to the same levels during the FMSOM phase. Here we used the signal captured at different temperatures during pyrolysis of SOM to evaluate how it relates to the amino sugars produced during the incubation (Figure 6). Overall, we observed that the content of amino sugars in the soil was negatively related to the signal captured at lower temperatures (< 400°C) and positively related to the signal captured at higher temperatures (> 400°C) with the exception of muramic acid. Further, we observed that fungal glucosamine measurements correlated to the SOM signal at the higher temperatures (450-550°C) (Fig 6a). Moreover, when evaluating the biological origin of the necromass (i.e. inoculation treatments) by the SOM thermal-signal it was the BF_complex_ treatment that generated the strongest thermal-stable signal (Fig. 6b). Despite F_only_ treatment showing the highest potential for necromass accumulation efficiency (Figure 4), the SOM generated here did not correlate to C measured during ramped thermal RE ramped pyrolysis (Figure 6b).

**Figure 6.**
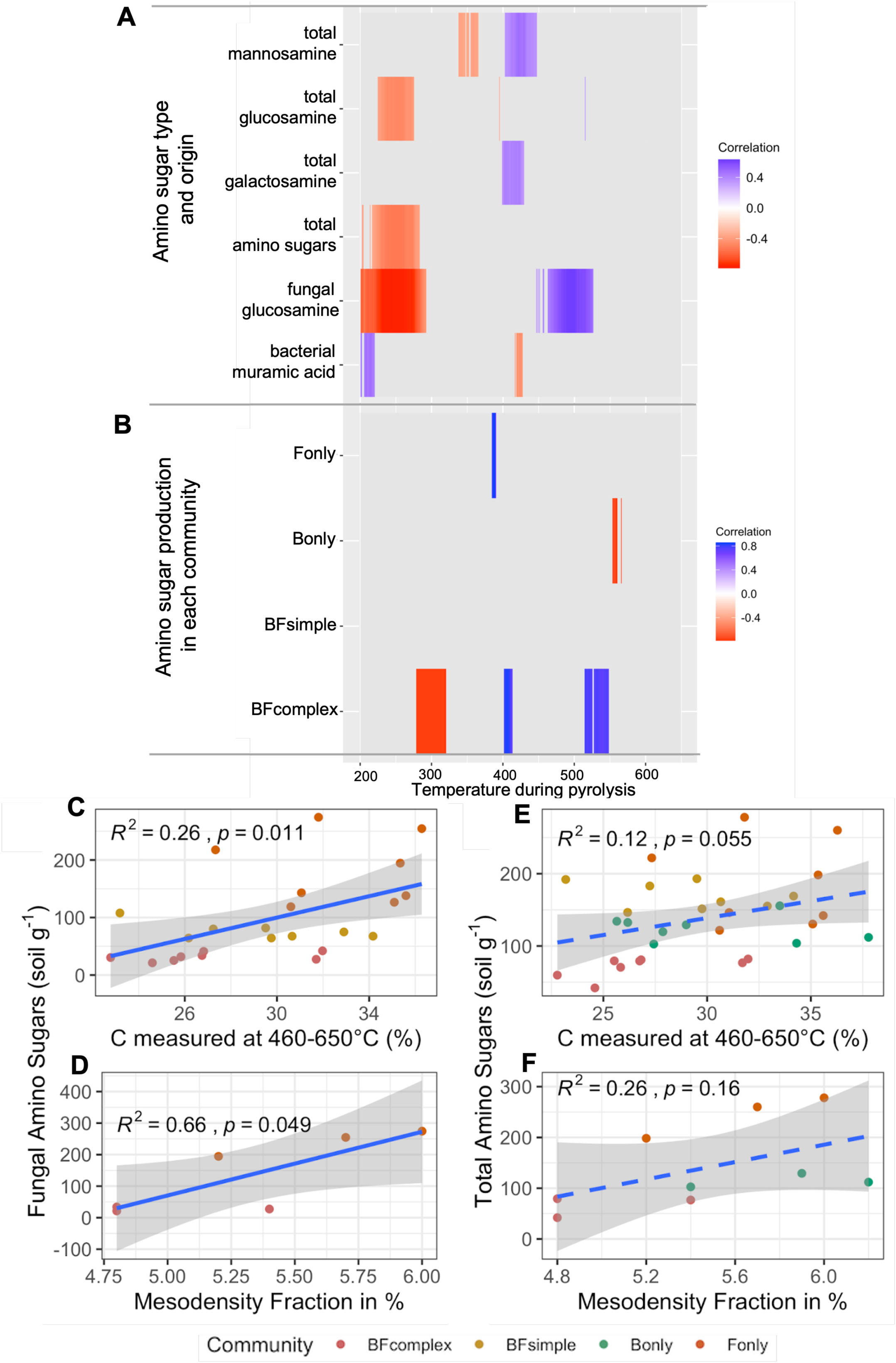
Relationship between the amino sugar residues, amino sugar biological origin according to incubation treatments and the SOM thermal-signal and the proportion of organo-mineral aggregate formed. Heat map of amino sugar residues and the FID signal captured during RE pyrolysis (A), relationship between amino sugars of fungal origin and the thermal-signal captured between 460-650 °C (B), relationship between fungal amino sugars or total amino sugars (fungal and bacterial origin) and the thermal-signal captured between 460-650 °C (C, E), and fungal amino sugars or total amino sugars and the mass proportion of meso-density aggregate formed (D, F).

### Evaluating the dynamics and persistence of SOM of distinct microbial origin during the second incubation

In the second phase of the experiment we sterilized the microcosms and inoculated all soils with an equivalent BFcomplex community from a natural agricultural soil (Figure 1) to evaluate how this new community responds to the distinct microbially-derived SOM formed during the FMSOM phase of the experiment. We did not observe significant differences in the organic carbon remaining in these soils at the beginning of the second incubation phase (Supplementary Figure 1). When evaluating the respiration, growth and CUE in the PSOM phase, we explained a more significant fraction of the variation in the respiration dataset compared to growth or CUE (79%, 44% and 14% for respiration, growth and CUE, respectively) (Figure 7). Moreover, we observed that amino sugars were negatively related to both respiration and growth measurements during the PSOM phase in these soils (Figure 7). We also evaluated the relationship between amino sugar accumulated in the first phase of the experiment and the measurements observed in the PSOM phase for each individual community type (Figure 8). The F_only_ community was the only treatment in which we did not observe any significant relationship between amino sugar produced during the FMSOM phase and cumulative respiration measured during 360 hours (15 days), short-term respiration during CUE incubations and growth measurements. In all other treatments we observed a negative relationship between total amino sugar and some of the activity measurements with the exception of CUE, which was not significantly related to amino-sugar content in any of the community types (Figure 8). As CUE is the compilation of both respiration and growth, the fact that CUE was not negatively correlated to amino sugar shows that the response of respiration and growth to amino sugar content was dissimilar.

**Figure 7.**
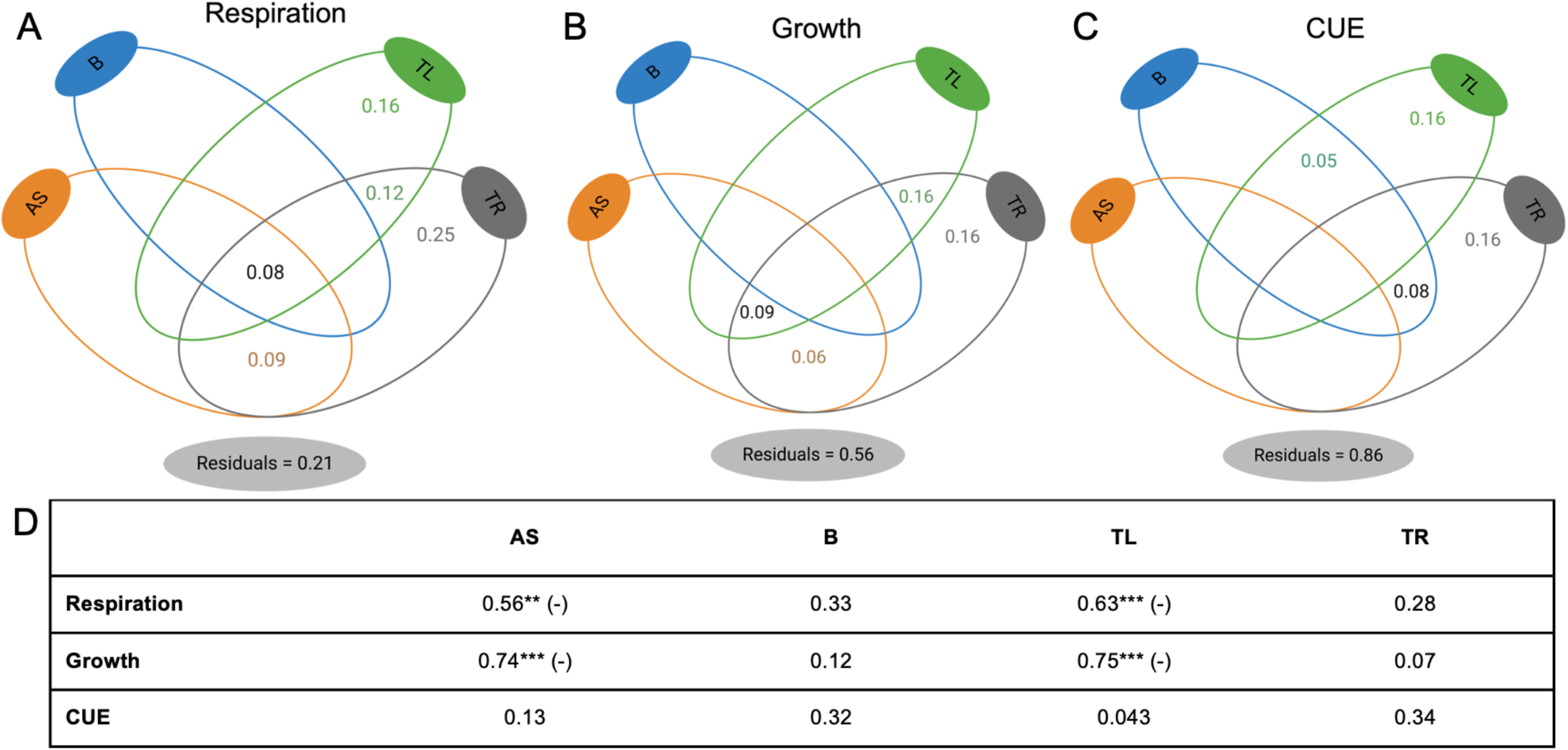
Variation partitioning of C cycling processes measured at PSOC in relation to the amino-sugar content, biomass and SOM thermal-signal measured at FMSOM. Variance was partitioned into amino sugar (AS), microbial biomass (B), SOC thermal labile signal (TL), SOC thermal resistant signal (TR) and by combinations of these predictors. Variance partitioning of respiration (a), growth (b) and CUE (c). Table with correlation coefficients in addition to the directions of the correlations in case of positive (+) or negative (−) if significant (* P < 0.05, **P < 0.01, ***P < 0.001) (D).

**Figure 8.**
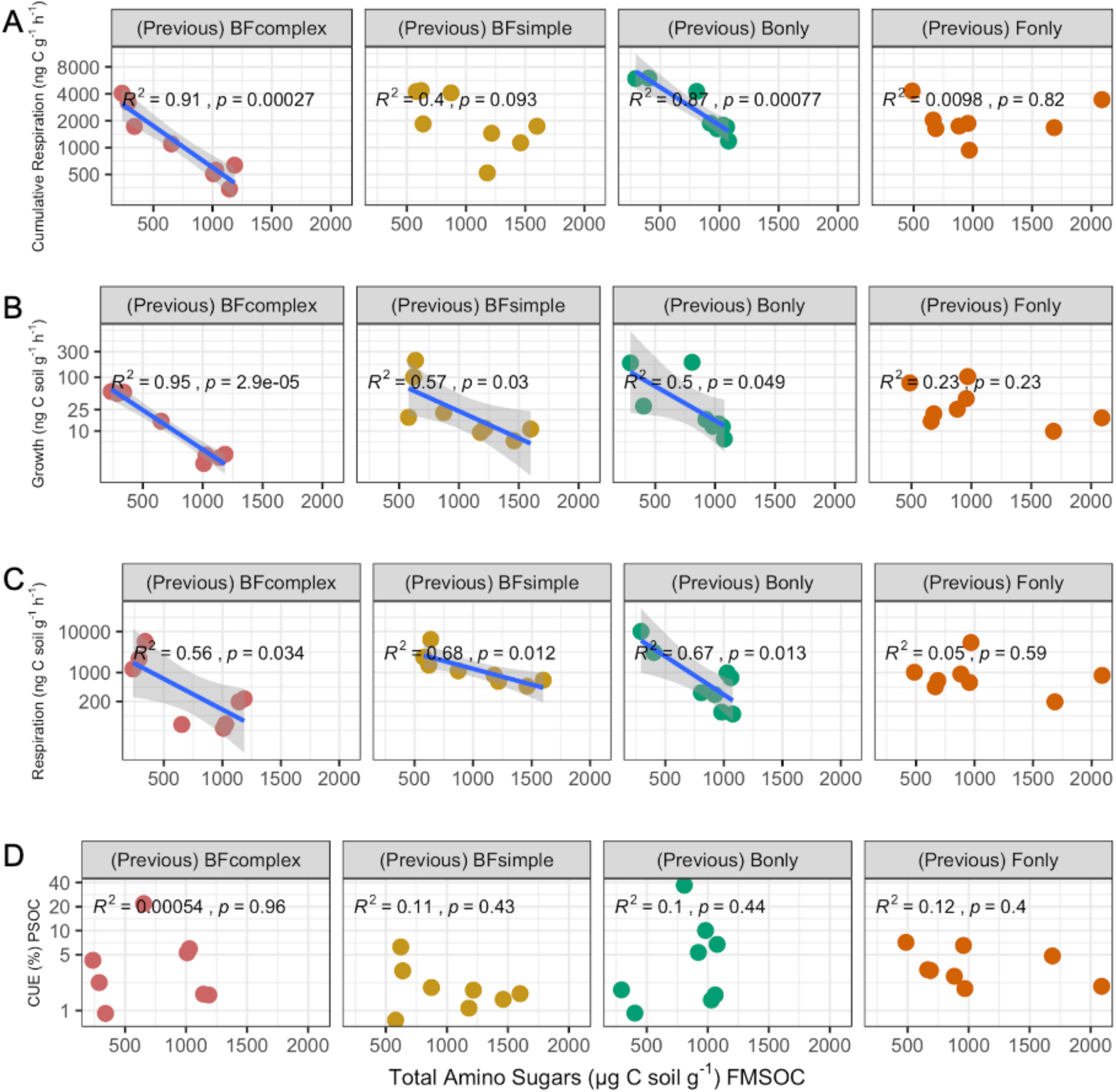
Relationship between total amino sugar measured at FMSOM and microbial C-cycling processes measured during PSOM for each community type. **A)** Relationship between the microbial activity measured as cumulative respiration for two weeks during PSOM phase and the amino sugar-C content. **B)** Relationship between growth measured at the end of PSOM and the amino sugar-C content. **C)** Relationship between respiration measurement at the end of PSOM and the amino sugar C content. **D)** Relationship between CUE measured at the end of PSOM and the amino sugar C content.

## Discussion

We manipulated microbial communities to control the necromass formation in model soils to test if necromass of different biological origins and formed under different abiotic conditions differ in its C thermal-signature and how this relates to its persistence in soils. We found evidence that fungal necromass is more thermally-stable than bacterial-necromass, when this necromass is formed under the occurrence of bacterial growth, suggesting the importance of microbial interactions for SOC stabilization. While we have an understanding that necromass is an important component of SOM^41,51^ varying from 33% in soil forest to up to 62% in grassland soils^10^, the role of necromass biological origin and dynamics within the microbial community for necromass persistence remains largely elusive. Our results show that while necromass chemistry was influenced by the communities, its total abundance was not, suggesting that the same amount of necromass (per unit of organic C) was formed in the distinct treatments. This allows us to evaluate the relative influence of necromass on SOM turnover and its influence on microbial carbon cycling processes in these soils^52^. To evaluate how the distinct biological origin and conditions of necromass formation would impact its persistence and SOM turnover, we analyzed the relationship between amino-sugars abundance and the FID signal captured during the pyrolysis phase of rock-eval. Overall, necromass content in the distinct treatments was positively related to the thermal-signal at higher temperatures and negatively related to the signal at lower temperatures, suggesting that microorganisms have consumed the labile substrate (captured at lower temperatures) and microbial growth and death (i.e. turnover) contributes to a more thermal-stable SOM signature in these soils. The only exception was the muramic acid of bacterial origin which showed a weak but positive relationship with the signal at lower temperatures and negative relationship with the signal captured at higher temperatures. These results suggest that necromass of solely bacterial origin is not very persistent in soil. Surprisingly, F_only_ treatment necromass signal was not related to the thermal signal captured at higher temperatures while the necromass measured at the BF_complex_ treatment was positively related to the signal at higher temperatures suggesting that concomitant growth of fungi and bacteria resulted in a more thermal-stable SOM signature. Buckeridge (2020) observed that necromass chemistry influences necromass-necromass (i.e. organic-organic) interactions resulting in higher retention of bacterial necromass if yeast necromass was present. Our experimental design does not allow us to differentiate between fungal x bacterial interactions of active microorganisms or dead organic-organic necromass interactions as hypothesized previously by Buckeridge *et al*. ^53^. Nevertheless, our results show that necromass measured under a more diverse microbial community (BF_complex_) is related to a more thermally-stable SOM signature compared to the BF_simple_ treatment, which corroborates to our previous findings showing that fungal and bacterial Simpson diversity index was positively correlated to a SOM chemical signature requiring higher energy for its thermal-decomposition^17^. In a previous experiment^17^, we observed that in the absence of fungi, bacterial communities produced a small pool of the extracellular enzyme to degrade chitin (i.e. N-acetylglucosaminidase) which is one of the components of the fungal cell wall and thus contributes to the necromass pool. Here we hypothesized that bacterial cells growing in the presence of fungi responded to the fungi presence by producing extracellular enzymes that resulted in further modifying and degrading the fungal cell wall which would increase biomass-turnover and the stabilization of the microbial-derived C. Accordingly, we observed the extracellular enzymatic pool to be the stronger driver of the SOM thermal-signature in a previous model soil experiment^17^. Alternatively, fungal biomass could also represent a source of organic nitrogen for community growth as N-limitation has been observed to be an important driver of microbial CUE^54^.

Necromass persistence is likely to be linked with organo-mineral aggregation based on the fact that micro and submicron-sized aggregates (also called organo-mineral complexes) isolated from bulk soil by various physical fractionation techniques are characterized by strong microbial signatures such as low C:N ratio, high delta 15N, amide-rich moiety, and by more negative Δ^14^C (less input of modern C)^42^. We observed a positive relationship between fungal-derived amino-sugars and the soil mass measured at the mesodensity fraction, suggesting the relevance of fungal necromass for soil aggregation, as observed previously^55^. The agglomeration of fine mineral particles promoted by fungal necromass drives microaggregate formation and soil structure development^31^. These results suggest that fungi necromass stimulates soil aggregation which is also known as mechanisms of OM occlusion and therefore protection against further decomposition within soil aggregates^32,42^. Accordingly, we observed a positive relationship between fungal amino-sugar content in these soils and SOC pyrolyzed within 460-650^0^C, while the total amino sugar content in these soils did not correlate to the more thermal stable C fraction (460-650^0^C) nor mesodensity fraction. These results corroborate our previous findings of enhanced soil aggregate formation with the presence of fungi in model soils^16^, suggesting that fungal necromass play a more important role than bacterial necromass for aggregate formation. Future studies should further evaluate to which extent soil aggregation influences the SOM thermal-signature and thermal-stability.

Previous studies have demonstrated that necromass accumulation efficiency depends on metabolic efficiency and other drivers such as clay content and mineralogy^41^. In this study our focus was on the influence of the microbial community, and therefore we used the same clay type and content across all soils. We observed that the F_only_ at high moisture treatment respired on average less than the other treatments, resulting in a higher amino-sugar accumulation efficiency compared to BF_complex_. We also conceived our experiment to evaluate if the distinct communities would exhibit contrasted carbon use efficiencies which could play a role in necromass formation. We showed that microbial physiology (i.e. respiration, growth and CUE) measurements performed at the end of FMSOM phase did not differ very strongly among community types or moisture treatments. Abiotic drivers such as moisture are considered important drivers of microbial activities, however in our system moisture had a limited effect on necromass formation and microbial activity parameters. This could be due to the fact that our lower moisture content did not exert physiological stress on the microbial communities. We did however observe a trend of higher microbial growth than respiration on more wet soils resulting in a tendency of higher CUE in wet compared to drier soils. Lower CUE in drier conditions has been previously observed in natural soils^29^and simple model soils^16^. Fungal dominated communities (F_only_), which are expected to be less sensitive to drier conditions^25,57^ showed to be less influenced by the two water moisture treatments, while communities containing bacteria showed a tendency to respond to moisture changes.

We observed that the necromass content formed in these soils was negatively related to respiration and growth suggesting that necromass constitutes a fraction of SOM not easily available to the microbial community re-inoculated in these soils as previously hypothesized^41,51,53^. While previous studies have highlighted the relevance of biotic-abiotic interactions for necromass retention (necromass-mineral interaction)^58^, our results highlight that biotic interactions may also play a role in necromass formation and persistence, as we observed that fungal necromass formed in the absence of bacterial growth (F_only_) was not negatively related to microbial activities in contrast to the necromass formed in all other communities. It has been recently hypothesized^59^ – with some empirical support – that biotic interactions are underestimated drivers of microbial growth efficiency^60,61^ which is a microbial trait that can influence microbial biomass and therefore necromass formation in soils in the long-term. We expected to observe a negative relationship between CUE and necromass content, however we did not observe such relationship during PSOM in any of the community treatments, although we recorded a negative relationship between amino-sugar content and both respiration and growth. While CUE is the composite measure of these other measures, we are hypothesizing that we did not observe a relationship with CUE because the catabolic and anabolic physiological responses of the microbial community to necromass presence in the soil were dissimilar. A possible explanation is that necromass content can differently impact growth and respiration if microorganisms are allocating more C to non-growth metabolites^11^ like extracellular enzymes in an attempt to acquire substrate in an environment with increasingly less available OM. This could suggest more “expensive growth” in response to physiological costs due to increases in the extracellular enzymatic pool and other costs related to acquisition of substrate and therefore we observe a de-coupling between respiration and growth as our measurement captures uniquely biomass increase^11,32^.

Altogether our results show that the biological origin of necromass and the conditions under which necromass is formed (e.g. under fungal-bacterial co-existence) play a role in necromass persistence in soils. Model soils can be used as valuable tools to help us understand the mechanisms driving soil functioning^16,17,41,62,63^, but our results cannot be extrapolated to natural soils without caution as microbial communities in soil are highly diverse and fungal and bacterial cells co-occur in large numbers. We also know that fungi are expected to dominate the first stages of plant biomass decomposition being followed by bacterial growth^20^. Our results showing the importance of co-occurrence of fungi and bacterial growth for necromass persistence could suggest the relevance of microbial succession for SOC stabilization in soils. Future studies should further investigate the role of microbial succession in SOM stabilization and whether there are potential context dependencies.

## Data and code availability

The data and R code supporting the findings presented here are available from the corresponding author on request and from the Open Science Framework Repository (https://osf.io/XXX).

### Autor contributions

L.A.D-H. and S.L. conceived the experiment. S.L., L.A.D-H., N.R. and V.L. conducted the experiments. The amino-sugar measurements were performed at the Key Laboratory of Vegetation and Environmental Change at the Chinese Academy of Sciences (Beijing, China) by X.L. under the supervision of X.F. The density fractionation was performed at the Institute for Agro-Environmental Sciences (Ibiraki, Japan) by R.W. D.B.N. performed the stable isotope analyses. S.L. analyzed the data under the supervision of L.A.D-H. All authors contributed to data interpretation. S.L. and L.A.D-H. wrote the first draft of the paper and all authors contributed to revising the manuscript.

## Acknowledgements

Funding for this project was provided by the Nessling Foundation and the Academy of Finland (STN MULTA; 327222) to A-L.L. This work was also conducted with support from the Zurich-Basel Plant Science Center fellowship to L.A.D-H., A-L.L. and A.K.

