## Supplementary material for "Who is who in necromass formation and stabilization in soil? Unraveling the role of fungi and bacteria as complementary players of biogeochemical functioning": Supplementary Material_Lepori et al..pdf

**Supplementary Table 1.** Composition of different communities. The single microbes were ordered from the catalogue of the DSMZ collection. (https://www.dsmz.de/collection/catalogue/microorganisms/catalogue).

| Community composition | Microbial inocula |
|---|---|
| $BF_{complex}$ | Natural soil sample |
| $BF_{simple}$ | *Streptomyces sp.* (DSM 687), *Microvirgula aerodenitrificans* (DSM 736), and *Trichoderma koningii* (DSM 63059) |
| $B_{only}$ | *Streptomyces sp.* and *Microvirgula aerodenitrificans* |
| $F_{only}$ | *Trichoderma koningii* |

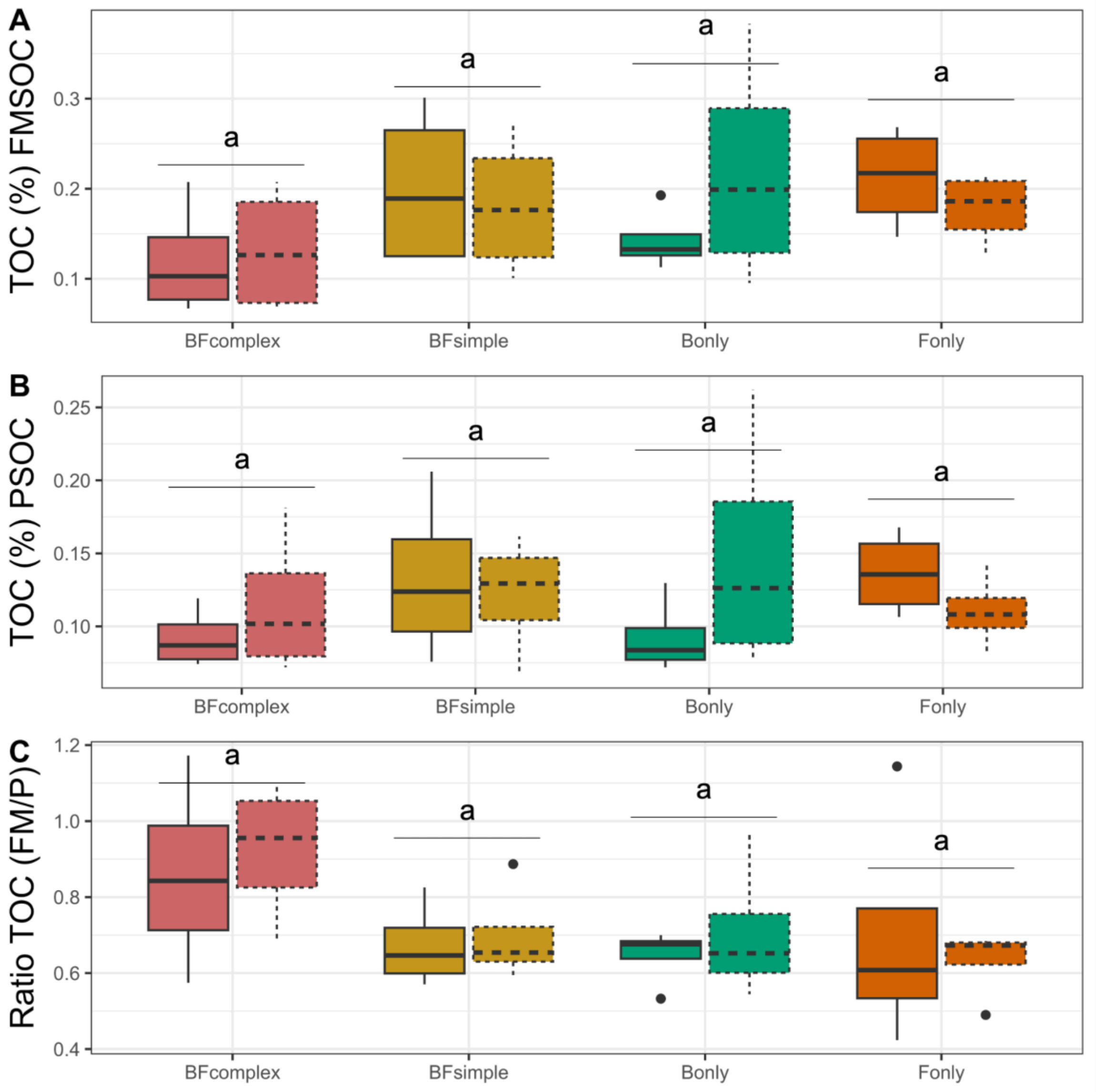


**Supplementary Figure 1. Total organic carbon (TOC) measured in these model soils.** TOC (%) measured at the end of the first incubation phase FMSOM (A), TOC (%) measured at the end of the second incubation phase PSOM (B) and the ratio between the TOC measured in the FMSOM and PSOM phases. Significant differences between treatments are indicated with different letters (anova followed by Tukey HSD test, $P < 0.05$) or significant t-test results ($P < 0.05$).

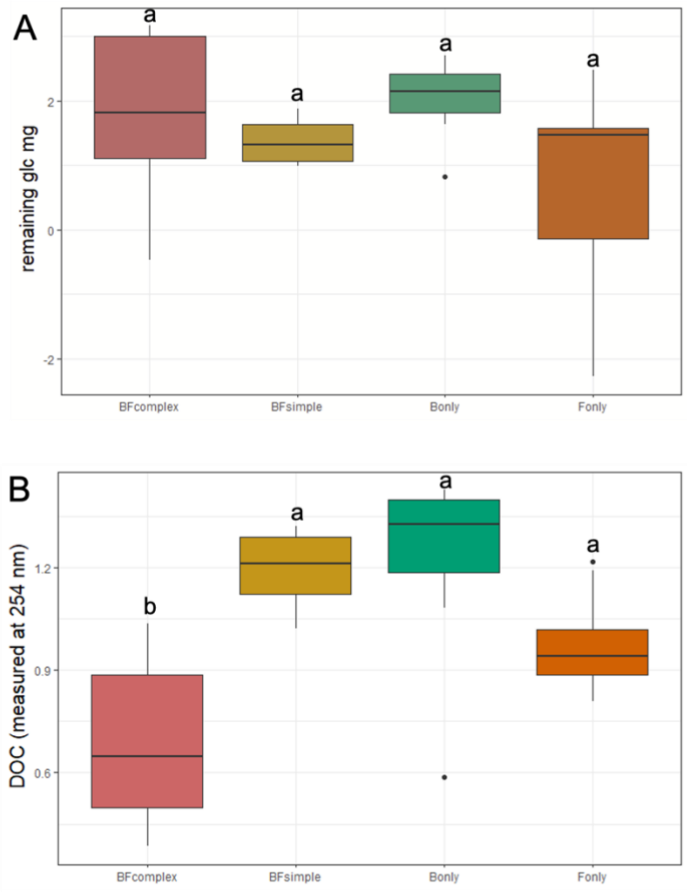


**Supplementary Figure 2. Estimates of organic carbon remaining in the soils at the end of FMSOM.** Estimated remaining glucose taking into account the cumulative respiration, necromass accumulated during the incubation and the microbial biomass measured at the end of FMSOC incubation (A). Absorbance measured at 254 nm as an estimate of how much dissolved organic carbon was present in the distinct soils. Significant differences between treatments are indicated with different letters (anova followed by Tukey HSD test, P < 0.05) or significant t-test results (*P* < 0.05).
